# Transcriptional and post-transcriptional mechanisms modulate cyclopropane fatty acid synthase through small RNAs in *Escherichia coli*

**DOI:** 10.1101/2024.02.12.579971

**Authors:** Colleen M. Bianco, Nancy N. Caballero-Rothar, Xiangqian Ma, Kristen R. Farley, Carin K. Vanderpool

## Abstract

The small RNA (sRNA) RydC strongly activates *cfa*, which encodes the cyclopropane fatty acid synthase. Previous work demonstrated that RydC activation of *cfa* increases conversion of unsaturated fatty acids to cyclopropanated fatty acids in membrane lipids and changes the biophysical properties of membranes, making cells more resistant to acid stress. The conditions and regulators that control RydC synthesis had not previously been identified. In this study, we demonstrate that RydC regulation of *cfa* is important for resistance to membrane-disrupting conditions. We identify a GntR-family transcription factor, YieP, that represses *rydC* transcription and show that YieP indirectly regulates *cfa* through RydC. YieP positively autoregulates its own transcription. We further identify additional sRNA regulatory inputs that contribute to control of RydC and *cfa*. The translation of *yieP* is repressed by the Fnr-dependent sRNA, FnrS, making FnrS an indirect activator of *rydC* and *cfa.* Conversely, RydC activity on *cfa* is antagonized by the OmpR-dependent sRNA OmrB. Altogether, this work illuminates a complex regulatory network involving transcriptional and post-transcriptional inputs that link control of membrane biophysical properties to multiple environmental signals.

**Importance:** Bacteria experience many environmental stresses that challenge their membrane integrity. To withstand these challenges, bacteria sense what stress is occurring and mount a response that protects membranes. Previous work documented the important roles of small RNA (sRNA) regulators in membrane stress responses. One sRNA, RydC, helps cells cope with membrane-disrupting stresses by promoting changes in the types of lipids incorporated into membranes. In this study, we identified a regulator, YieP, that controls when RydC is produced, and additional sRNA regulators that modulate YieP levels and RydC activity. These findings illuminate a complex regulatory network that helps bacteria sense and respond to membrane stress.

## Introduction

The gram-negative bacterial cell envelope confers protection from a variety of toxic agents, many of which are encountered by *Escherichia coli* and other enteric bacteria in their natural environments, such as solvents, acid, detergents, bile salts and antibiotics [1]. The envelope is also important for cellular responses to external stresses with alterations in membrane lipid and membrane protein composition playing critical roles in maintaining physiological homeostasis. Small RNAs (sRNAs) are well-characterized regulators that participate in maintaining cell envelope homeostasis through modulation of membrane protein synthesis [2–7]. Several sRNAs are integral parts of important stress regulons, including the α^E^, Cpx, and OmpR responses. The transcription factor σ^E^ maintains outer membrane homeostasis by inducing transcription of ∼100 genes, comprised mainly of genes encoding proteins involved in membrane repair as well as three sRNAs, RybB, MicA and MicL [3–5]. RybB and MicA down-regulate synthesis of abundant membrane porins, including OmpA, OmpC, OmpD, and OmpF, upon entry into stationary phase [4, 5, 8, 9]. MicL represses translation of outer membrane lipoprotein Lpp, a highly abundant protein [3]. Together MicA, RybB, and MicL repress protein synthesis of all the most abundant outer membrane proteins in response to stress [3]. While σ^E^ is responsible for maintaining homeostasis of the outer membrane, the Cpx response monitors inner membrane integrity [10]. The Cpx response controls production of an sRNA called CpxQ, which is processed from the 3’ UTR of *cpxP* mRNA [2]. Synthesis of CpxQ is triggered by membrane damaging stresses such as excessive synthesis of membrane proteins, alkaline pH, ethanol, and cationic antimicrobial peptides [2]. Upon induction, CpxQ represses the production of numerous inner membrane proteins [2]. Both CpxQ and the Cpx pathway are required for cell survival during conditions that lead to dissipation of membrane potential [2]. Lastly, the sRNAs OmrA and OmrB are induced by high osmolarity and are regulated by the OmpR response regulator [6]. Most mRNA targets are shared between OmrA and OmrB and encode outer membrane proteins. For example, OmrA and OmrB repress translation of *ompT*, which encodes an outer membrane protease, along with *cirA*, *fecA* and *fepA,* all encoding different outer membrane porins specific for iron–siderophore complexes [6].

Importantly, while changes in the structure of the fatty acids in phospholipids can alter critical membrane properties such as fluidity and permeability [1], the roles of sRNAs in modulating membrane lipids are just beginning to be explored. RydC is the first sRNA shown to regulate membrane fatty acid composition and plays a role in alteration of cell membrane properties and resistance to stress [11, 12]. RydC is a 65-nt sRNA that requires the RNA chaperone Hfq for stability and base-pairing interactions with target mRNAs [11, 13–15]. Like other Hfq-dependent sRNAs, RydC base pairing with target mRNAs has different outcomes depending on the site of interaction. RydC’s best characterized target is *cfa* mRNA, which codes for cyclopropane fatty acid (CFA) synthase, an enzyme that converts unsaturated fatty acids to cyclopropanated fatty acids [11, 12, 16]. RydC base pairs at a site within the long 5’ untranslated region (UTR) of *cfa* mRNA and modulates RNase E-dependent degradation and Rho-dependent transcription termination [11, 17]. We showed previously that RydC-mediated activation of *cfa* promotes survival during acid stress [12]. In addition to regulating fatty acid modification through activation of *cfa*, RydC has also been shown to impact mRNAs involved in sugar utilization (*yejABEF*), biofilm formation (*csgD*), and aromatic amino acid biosynthesis (*pheA* and *trpE*) [13, 15, 18].

Previous work identified two additional sRNAs, ArrS and CpxQ, that also directly regulate *cfa* [12]. Like RydC, ArrS acts through reducing *cfa* mRNA turnover leading to increased translation of *cfa* mRNA [12]. In contrast, CpxQ binds *cfa* mRNA upstream from the site bound by activating sRNAs. Together with Hfq, CpxQ binding may modulate susceptibility to RNase E cleavage site protected by the activating sRNAs. We recently showed that like RydC, ArrS and CpxQ also regulate *cfa* through modulation of Rho-dependent transcription termination in the long *cfa* 5’ UTR [17]. While the direct signals responsible for promoting sRNA regulation of *cfa* are not currently known, both CpxQ and ArrS were previously linked to cell envelope [2] and acid stress responses [19], respectively.

To better understand the physiological function of RydC and its role in cellular homeostatic or stress response pathways through *cfa* regulation, we sought to identify environmental signals and regulators that control RydC production. We found that RydC and *cfa* promote viability in the presence of solvents and other molecules that disrupt membrane integrity. A candidate-based approach to identify regulators revealed that none of the known membrane stress-responsive transcription factors regulate *rydC* transcription. Instead, a transposon screen identified a GntR-like regulator, YieP, as a repressor of RydC. We showed that YieP auto-activates its own transcription and that the sRNA regulator FnrS represses *yieP* mRNA translation. Additionally, we demonstrate that the osmotic stress-responsive sRNA OmrB inhibits RydC activity on *cfa*, suggesting that OmrB acts as a sponge for RydC. Collectively, these results demonstrate that RydC is regulated directly at both transcriptional and post-transcriptional levels within a regulatory network associated with membrane stress signals.

## Results

### Conservation of *rydC* genomic location and promoter region

RydC is a ∼65 nt sRNA that is conserved in several related enterobacterial species [11, 13]. The transcription of *rydC* is controlled by a σ^70^-dependent promoter with a perfect -10 box that matches the consensus sequence of ‘TATAAT’ in all species except *C. koseri* (Fig. 1A), but there is no obvious -35 box and there is little conservation upstream of the core promoter (Fig. 1A). A search for known or predicted transcription factor binding sites compiled in the *E. coli* Regulon Data Base did not reveal any sites within the *rydC* promoter [20].

**Figure 1.**
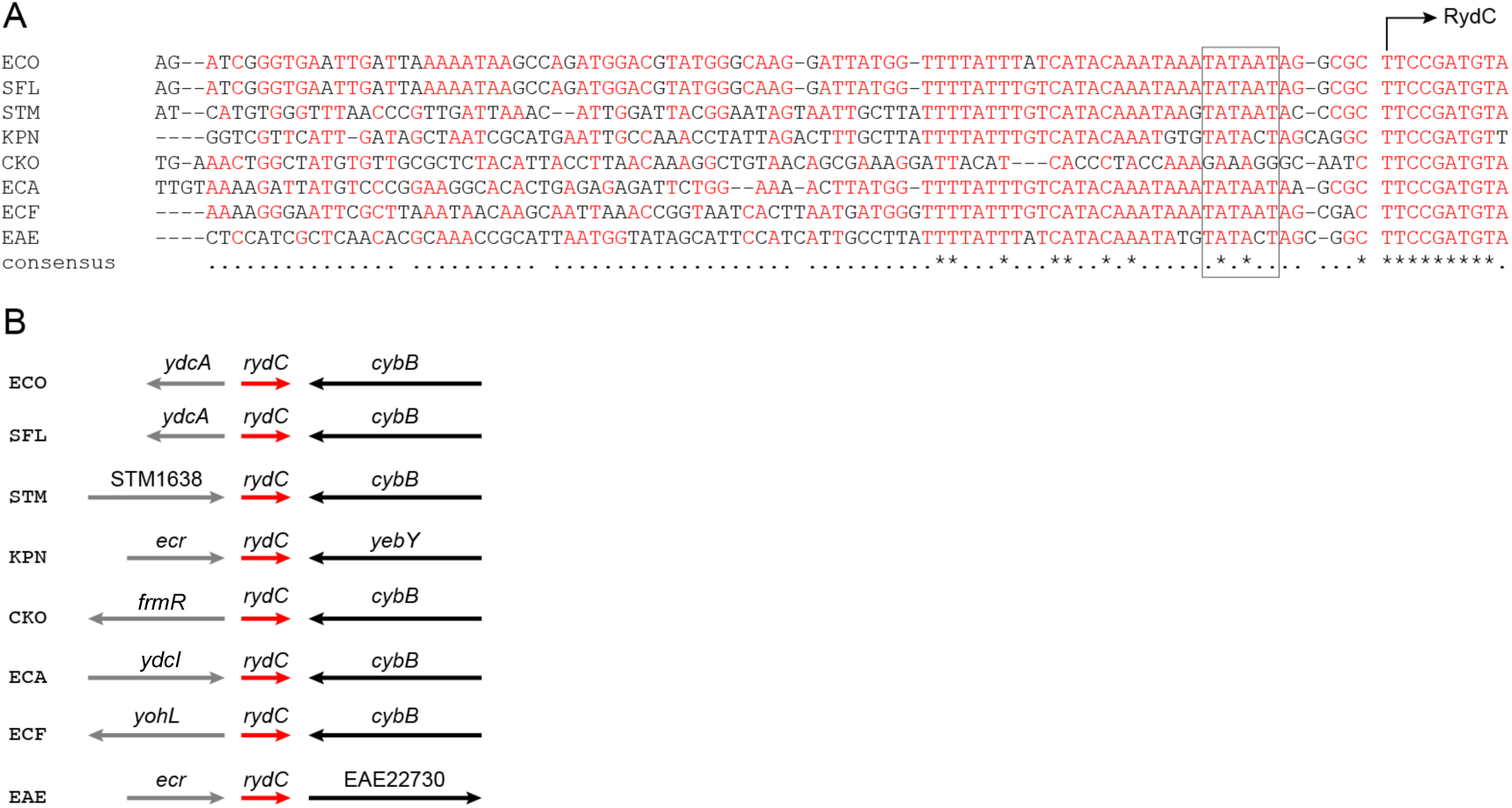
*rydC* promoter sequence and genomic context. (A) *rydC* promoter conservation. Sequences from eight enteric bacterial species were aligned using Clustal Omega. Species abbreviations: ECO, *Escherichia coli* str. K-12 substr. MG1655; SFL, *Shigella flexneri* 2a str. 301; STM, *Salmonella enterica* serovar Typhimurium LT2; KPN, *Klebsiella pneumoniae* HS11286; CKO, *Citrobacter koseri* ATCC_BAA_895; ECA, *Escherichia albertii* strain 1551-2; ECF, *Escherichia fergusonii* ATCC 35469; EAE, *Enterobacter aerogenes* KCTC 2190. The *rydC* transcription start site is marked with an arrow and the -10 element is indicated with a box. (B) *rydC* genomic location conservation. Species abbreviations are as in A. The *rydC* gene is a red arrow. Flanking genes are described in the text.

As it is common for genes encoding transcriptional regulators to be found in proximity to the genes they regulate, we looked at the chromosomal region surrounding *rydC* to identify putative regulators (Fig. 1B). In most species where *rydC* is found, the 3’ flanking gene is *cybB*, encoding cytochrome b_561_. In *E. coli* and *Shigella sp.* the 5’ flanking gene is *ydcA*, which encodes a small protein (57 aa) of unknown function. YdcA has a signal sequence and is predicted to be a periplasmic protein. In *Salmonella*, the *rydC* gene is in the intergenic region between STM1638, coding for a putative SAM-dependent methyltransferase, and *cybB.* In *Citrobacter koseri*, *rydC* is located next to *frmR*, coding for an FrmR/RcnR family transcriptional repressor. In *Escherichia fergusonii, yohL* coding for another FrmR/RcnR-like family transcriptional regulator, is found at the 5’ end of RydC. In *Escherichia albertii,* the gene RS00345, coding for a LysR family transcriptional regulator that is 98% similar to YdcI of *E. coli* flanks the 5’ end of *rydC.* In *Enterobacter aerogenes*, the *rydC* gene is located between Ecr encoding a hypothetical small membrane protein and EAE22730 encoding a spore coat domain-containing protein. EAE22730 is only found in *E. aerogenes* and a BLAST search did not reveal any homologues in *E. coli*. In *Klebsiella pneumoniae*, the *rydC* gene is located between Ecr and *yebY*, encoding a protein of unknown function.

To test whether RcnR, FrmR, or YdcI homologs regulate *rydC* transcription in *E. coli,* we compared activity of a *rydC*′-*lacZ* transcriptional fusion in wild-type (WT), *rcnR, frmR,* and *ydcI* mutant strains. There were no differences in fusion activity between these four strains (Fig. S1), suggesting that *E. coli rydC* transcription is not controlled by regulators homologous to those encoded by genes co-localized with *rydC* in related enterobacterial species.

### RydC and Cfa play a role in resistance to membrane stresses

To investigate conditions relevant to regulation of *rydC,* we examined RydC-dependent phenotypes. Because RydC alters membrane composition by increasing CFA levels, we tested the role of RydC in promoting resistance to butanol, a treatment shown to induce CFA formation [21] (Fig. 2A). WT and Δ*rydC* mutant strains were grown on LB plates containing 1% butanol overnight at 37 °C. The WT strain carrying vector control, P_lac_-*rydC* or P_lac_-*cfa* plasmids grew well on butanol (Fig. 2A, left). In contrast, the Δ*rydC* strain had significantly impaired growth (Fig. 2A, Δ*rydC* P_lac_-vector, right). The sensitivity of the Δ*rydC* strain to butanol stress was partially reversed by P_lac_-*rydC* (Fig. 2A, left) and fully reversed by P_lac_-*cfa* (Fig. 2A, right) plasmids. This suggests that the growth defect of the Δ*rydC* strain on butanol is primarily due to the inability to activate *cfa*. Consistent with this hypothesis, the Δ*cfa* strain also displayed a growth defect in the presence of butanol (Fig. 2A, right). This defect was restored by complementation with P_lac_-*cfa* (Fig. 2A, left). Importantly, production of RydC in the *cfa* mutant background did not restore growth on butanol (Fig. 2A, right). Together, these results suggest that RydC is an important activator of *cfa* during stress caused by butanol. Furthermore, the activation of *cfa* by RydC appears to be a key determinant of cell growth under these conditions.

**Figure 2.**
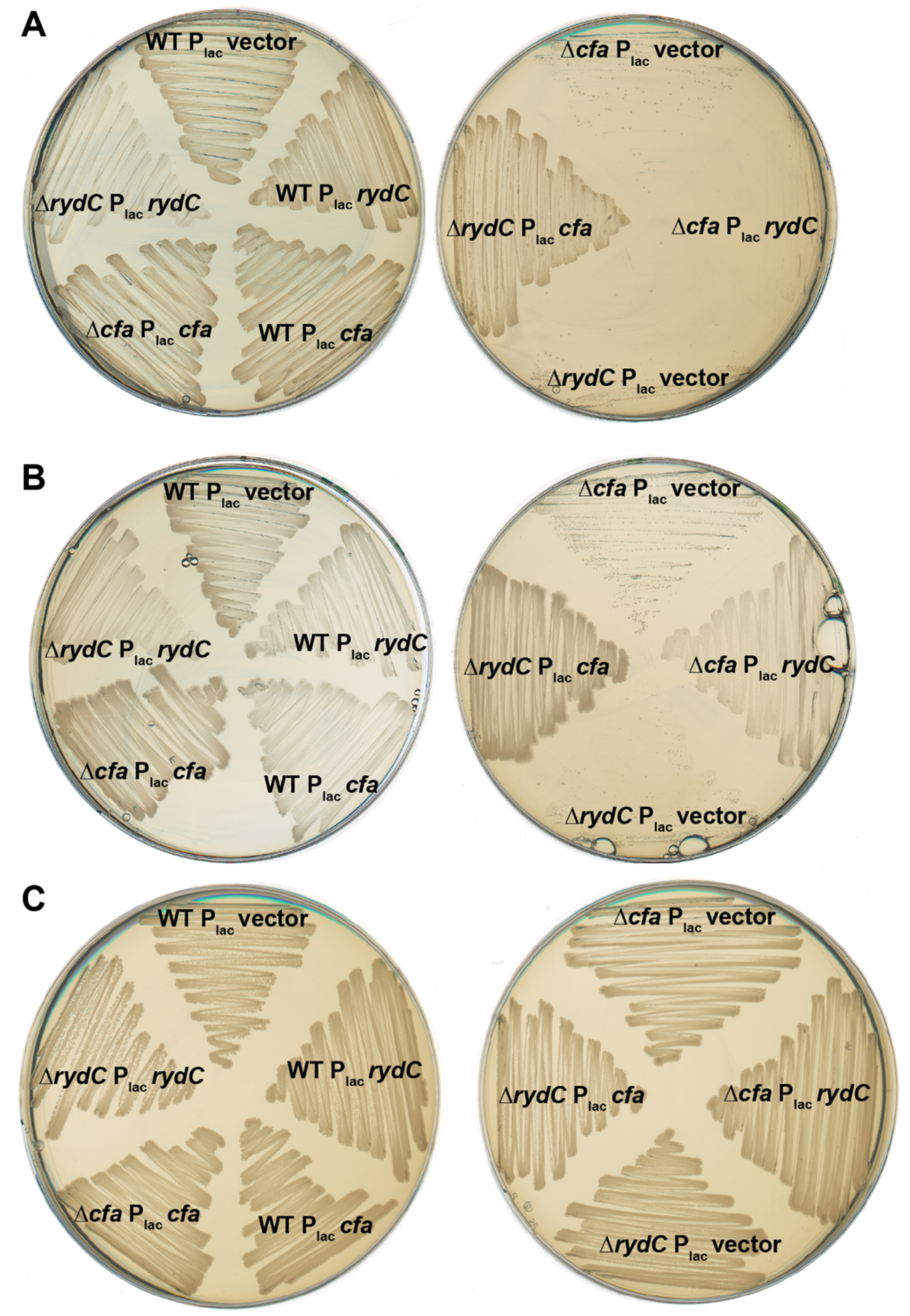
Phenotypes due to cell membrane stress. *E. coli* strains with the indicated plasmids were plated on LB containing 1% v/v 1-butanol (A), 0.5% v/v SDS and 0.5 mM EDTA (B) or no additions (C). All plates contained ampicillin and IPTG and were incubated overnight at 37°C.

We next tested if the Δ*rydC* mutant was sensitive to other compounds that perturb the cell envelope, such as sodium dodecyl sulfate (SDS) and ethylenediaminetetraacetic acid (EDTA). We evaluated growth of WT and Δ*rydC* mutant strains on LB containing 0.5% SDS and 0.5 mM EDTA. Growth of the Δ*rydC* strain (Fig. 2B, right) was reduced compared to the WT strain (Fig. 2B, left). Growth of the Δ*rydC* mutant was restored when P_lac_-*rydC* (Fig. 2B, left) or P_lac_-*cfa* (Fig. 2B, right) was supplied on a plasmid. The Δ*cfa* strain was slightly inhibited by SDS/EDTA (Fig. 2B, right) but was complemented by P_lac_-*cfa* (Fig. 2B, left). Interestingly, in contrast with the butanol phenotype, the Δ*rydC* mutant is more sensitive to SDS/EDTA compared to the Δ*cfa* mutant. These observations suggest that RydC activation of *cfa* is important for resistance to stress caused by SDS/EDTA, but that RydC regulation of targets in addition to *cfa* may play a role in reducing cell envelope stress caused by these compounds.

### Transcription of *rydC* does not respond to known membrane stress conditions and regulators

We monitored activity of a *rydC*′-*lacZ* fusion in the absence and presence of SDS/EDTA and butanol and other compounds known to induce membrane stress. Even though we see pronounced growth phenotypes for *rydC* mutants growing in the presence of SDS/EDTA or butanol, these compounds did not alter activity of the *rydC*′-*lacZ* fusion compared to the control condition (Fig. S2A, B). Other membrane stress inducing compounds and conditions including indole, bile salts, lidocaine, low pH, ethanol, imidazole, polymyxin B, and CCCP also did not alter activity of the *rydC*′-*lacZ* fusion compared to the control (Fig. S2A, B).

To test whether any known envelope stress response transcription factors play a role in modulating *rydC* transcription, we tested activity of a *rydC*′-*lacZ* transcriptional fusion in WT and mutant strains lacking known cell envelope stress regulators. Mutants lacking the BaeSR two-component system, the global stress response regulator σ^S^ (encoded by *rpoS*), and the Rcs signal transduction system had similar levels of *rydC* promoter activity as the wild-type strain (Fig. 3A). To assess the role of the σ^E^-dependent envelope stress response, we tested *rydC*′-*lacZ* activity in an *rseA* mutant, which lacks the anti-σ factor that normally keeps σ^E^ inactive. (An *rpoE* null mutant is inviable because σ^E^ is essential in *E. coli.*) The *rseA* mutant strain and WT strain also had similar levels of *rydC*′-*lacZ* (Fig. 3A). Mutation of genes encoding components of the phage shock protein (*psp*) system induced *rydC* transcription slightly, but the effect was small (less than 2-fold compared to the wild-type strain, Fig. 3A). We next tested regulators involved in more specific stress responses, including fatty acid degradation (FadR), the global transcription factor Lrp, iron uptake (Fur), carbon flow (Cra), osmotic stress (OmpR), aerobic respiration control (ArcB), NAD synthesis and transport (NadR) and control of pyruvate dehydrogenase (PdhR). Mutant strains lacking most of these regulators had similar levels of *rydC*′-*lacZ* activity compared to the wild-type strain (Fig. 3B). The *lrp, fadR,* and *pdhR* mutant strains had slightly reduced transcription of *rydC*′-*lacZ*, but again the effects were modest (Fig. 3B). The CpxAR two component system responds to stresses that disrupt the inner membrane and alter redox balance [10, 22, 23]. To test whether CpxAR regulates *rydC* transcription, we compared *rydC*′-*lacZ* activity in WT, Δ*cpxR,* and Δ*cpxA* mutant backgrounds and found no difference compared to the WT strain (Fig. 3C). Collectively, these data suggest that *rydC* transcription is not responsive or minimally responsive to known envelope stress response regulators and other characterized stress response pathways.

**Figure 3.**
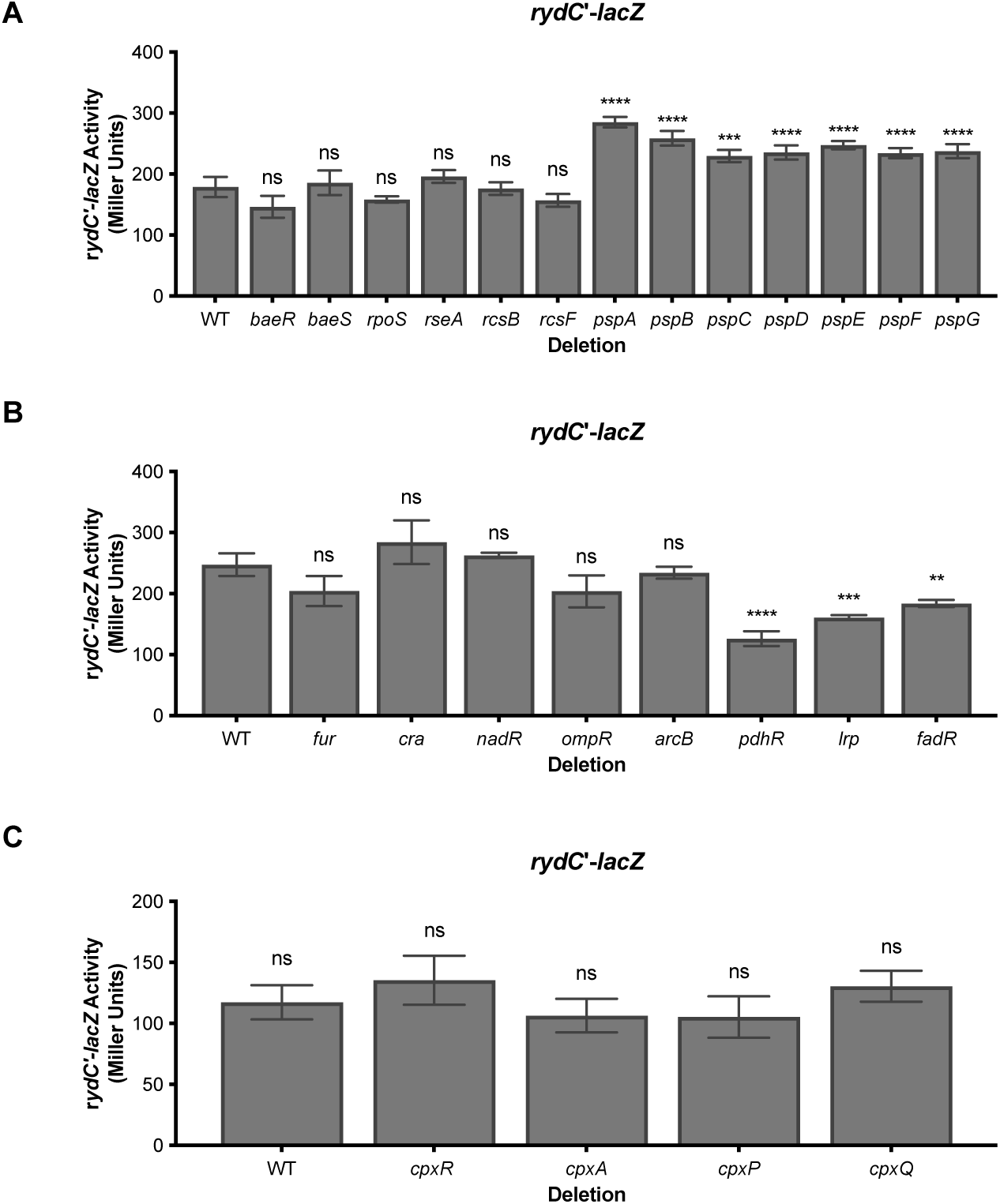
Testing candidate membrane stress regulators for a role in *rydC* transcription. Strains harboring a *rydC’-lacZ* transcriptional fusion with deletions of indicated candidate regulators were constructed as described in Materials and Methods. Strains were grown overnight in TB then subcultured 1:100 to fresh TB medium. Subcultures were grown at 37°C with shaking for 3 hours and then β-galactosidase assays were performed. Error bars are standard deviations of the results of three independent experiments. Statistical significance was determined using a one-way ANOVA followed by Tukey’s post hoc test. Only significant differences between activity of WT compared to mutants are shown (ns, P not significant; *, P < 0.05; **, P < 0.01; ***, P < 0.0001; ****, P < 0.0001).

### GntR-family regulator YieP represses transcription of *rydC*

Since a candidate-based approach did not reveal any known transcription factors participating in *rydC* regulation, we next performed transposon mutagenesis on a strain carrying a *rydC*′*-lacZ* transcriptional fusion. We isolated mutants with transposon insertions in the *yieP* gene that had strongly increased *rydC*′*-lacZ* activity. YieP is a helix-turn-helix-type transcriptional regulator that has recently been implicated in 3-hydroxypropionate tolerance due to its regulation of genes encoding the transporter YohJK [24, 25]. We validated YieP-dependent regulation of *rydC* transcription in both *E. coli* and *Salmonella* (Fig. 4). Each WT or Δ*yieP* host carried a *rydC*′*-lacZ* transcriptional fusion and either a vector control or plasmid with an inducible copy of *yieP.* Deletion of *yieP* resulted in a 20-fold increase in *rydC*′*-lacZ* activity in *E. coli* (Fig. 4A), and a 7.5-fold increase in *Salmonella* (Fig. 4B). Complementation with plasmid-borne *yieP* reduced *rydC*′*-lacZ* activity in both hosts (Fig. 4A, B). We performed Northern blots probing for RydC on samples harvested from WT and Δ*yieP E. coli* strains (Fig. 4C). RydC was produced at detectable levels only in the Δ*yieP* strain and the positive control strain where RydC was expressed ectopically (Fig. 4C). These experiments confirmed the results from the transcriptional reporter fusions and suggest that YieP acts as a repressor of *rydC* transcription.

**Figure 4.**
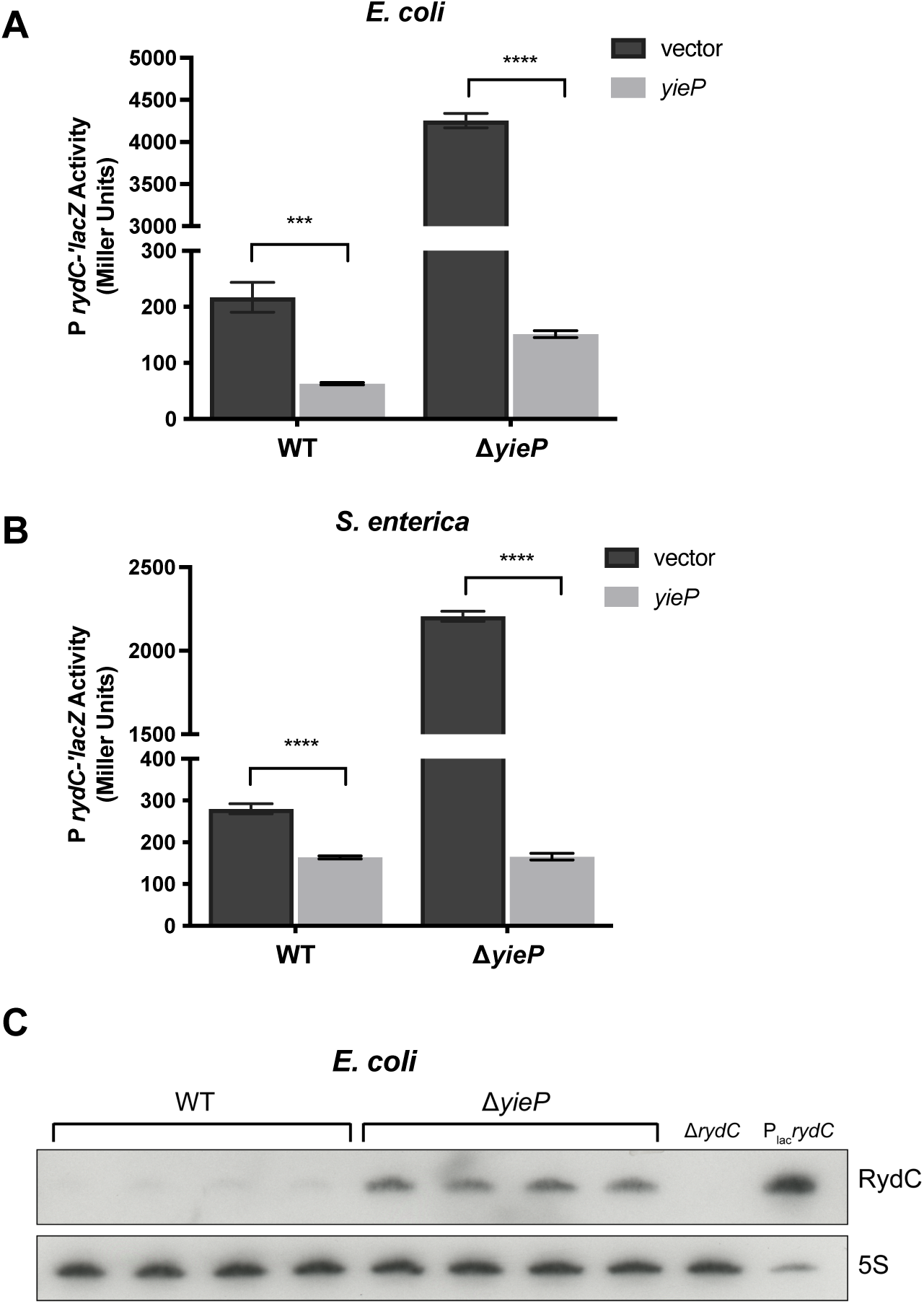
YieP represses *rydC* transcription in *E. coli* and *Salmonella*. β-galactosidase activity was measured in wild-type (WT) and Δ*yieP* strains containing a *rydC*′-*lacZ* transcriptional fusion with either a vector or P_lac_-*yieP* plasmids in *E. coli* (A) or *S. enterica* serovar Typhimurium (B). Strains were grown overnight with Amp and 0.1 mM IPTG then subcultured 1:100 to fresh TB medium containing Amp and 0.1 mM IPTG. Subcultures were grown at 37°C with shaking for 3 hours and then β-galactosidase assays were performed. All experiments were performed at least in triplicate in three independent replications. Error bars are standard deviations of the results of three independent experiments. Statistical significance was calculated by using a two-tailed unpaired t-test, (*, P < 0.05; **, P < 0.01; ***, P < 0.0001; ****, P < 0.0001). (C) RydC levels in WT or Δ*yieP E. coli* cells measured by Northern blot. Cells were grown overnight then subcultured 1:100 to fresh LB medium (four biological replicates). Subcultures were grown for 2 hours and then RNA was extracted. A Δ*rydC* strain served as a negative control. 5S RNA served as loading control.

The *yieP* gene is conserved in *E. coli* and *Salmonella*, as well as other Enterobacteriaceae, and is found in a conserved genomic region (Fig. 5A). In every genome examined except that of *Escherichia fergusonii*, *yieP* is found in an operon with a gene encoding an uncharacterized Major Facilitator Superfamily (MFS) transporter (called HsrA in *E. coli*) (Fig. 5A). The *yieP* operon is found downstream of a 16S rRNA gene and upstream of an operon involved in ribose catabolism and transport (Fig. 5A). The amino acid sequence of YieP is also well conserved, especially the amino acid residues 40-63 which comprise the helix-turn-helix DNA binding domain (Fig. 5B). We used AlphaFold to predict the 3D structure of YieP (Fig. 5C) [26, 27]. The AlphaFold structure confirmed the nine alpha helices predicted from the YieP amino acid structure with high (>90%) confidence. The predicted structure shows the first 66 amino acids of the N-terminus, which includes the DNA binding domain, forms an ordered structure that is linked by n1 and n2 to the C-terminus portion of the protein, which contains the signal recognition domain. YieP is a member of the GntR family of proteins, so YieP most likely forms a homodimer, a hallmark of GntR-like regulators.

**Figure 5.**
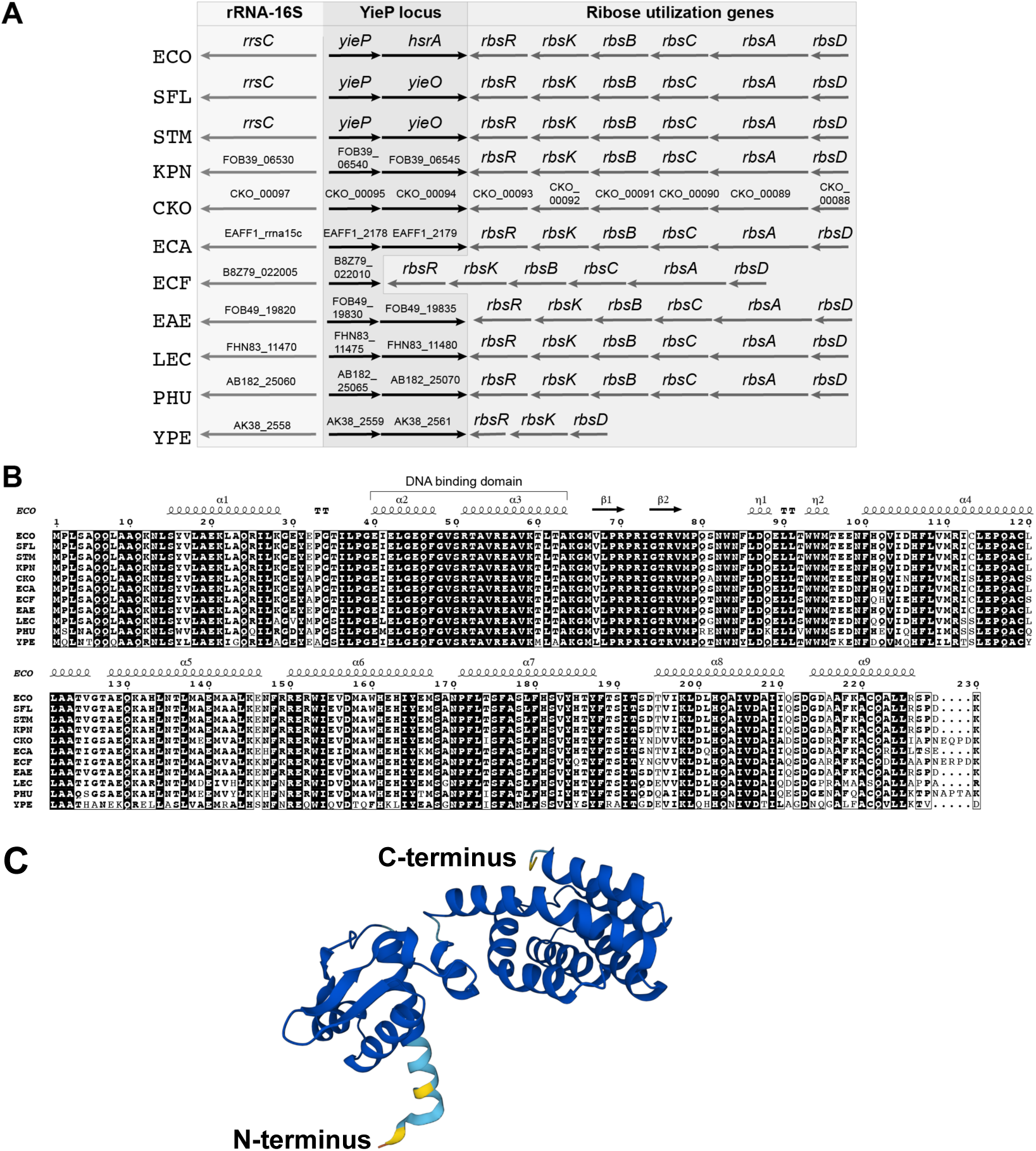
Characterization of YieP. (A) The genomic region of *yieP* is highly conserved. Strain abbreviations: ECO, *Escherichia coli*; SFL, *Shigella flexneri*; STM, *Salmonella enterica* serovar Typhimurium; KPN, *Klebsiella pneumoniae* (FDAARGOS_631); CKO, *Citrobacter koseri* (ATCC BAA-895); ECA, *Escherichia albertii* (KF1); ECF, *Escherichia fergusonii* (EFCF056); EAE, *Enterobacter aerogenes* (FDAARGOS_641); LEC, *Leclercia adecarboxylata* (Z96-1); PHU, *Phytobacter ursingii* (CAV1151); YPE, *Yersinia pestis* (CO92). (B) Amino acid sequence conservation of YieP. The predicted structure of YieP is labeled above the sequence and the highly conserved helix-turn-helix DNA binding domain is labeled. (C) Alpha fold structure of monomeric YieP in *E. coli* K12. Residues are colored by model confidence in the form of a per-residue confidence score (pLDDT) between 0 and 100: dark blue: very high (pLDDT > 90), light blue: confident (90 > pLDDT > 70), yellow: low (70 > pLDDT > 50).

We tested whether deletion of *yieP* affected RydC target gene expression. To date, three direct mRNA targets of RydC have been confirmed: *cfa* [11], *trpE* [18], and *pheA* [18]. RydC-mediated repression of *trpE* and *pheA* is modest, whereas RydC-mediated activation of *cfa* is substantial [18]. We constructed reporter strains carrying a previously characterized *cfa’-’lacZ* translational fusion [18]. As observed previously, Δ*rydC* strains have reduced *cfa’-’lacZ* activity compared to the WT parent, and RydC-producing strains have ∼9-fold increased activity (Fig. 6) because RydC stabilizes *cfa* mRNA and inhibits Rho-dependent termination through a base pairing-dependent mechanism [11, 17]. Activity of the *cfa’-’lacZ* fusion in the Δ*yieP* strain was ∼7-fold higher than in the WT background (Fig. 6). To determine whether this increase in activity was RydC-dependent, we measured *cfa’-’lacZ* activity in a Δ*yieP* Δ*rydC* strain and observed that activity in this strain was similar to activity in the Δ*rydC* strain. These results are consistent with the model that YieP represses *rydC* and prevents RydC production and subsequent activation of *cfa* in these growth conditions. Deletion of *yieP* leads to increased RydC levels (Fig. 4C) and activation of *cfa’-’lacZ* (Fig. 6).

**Figure 6.**
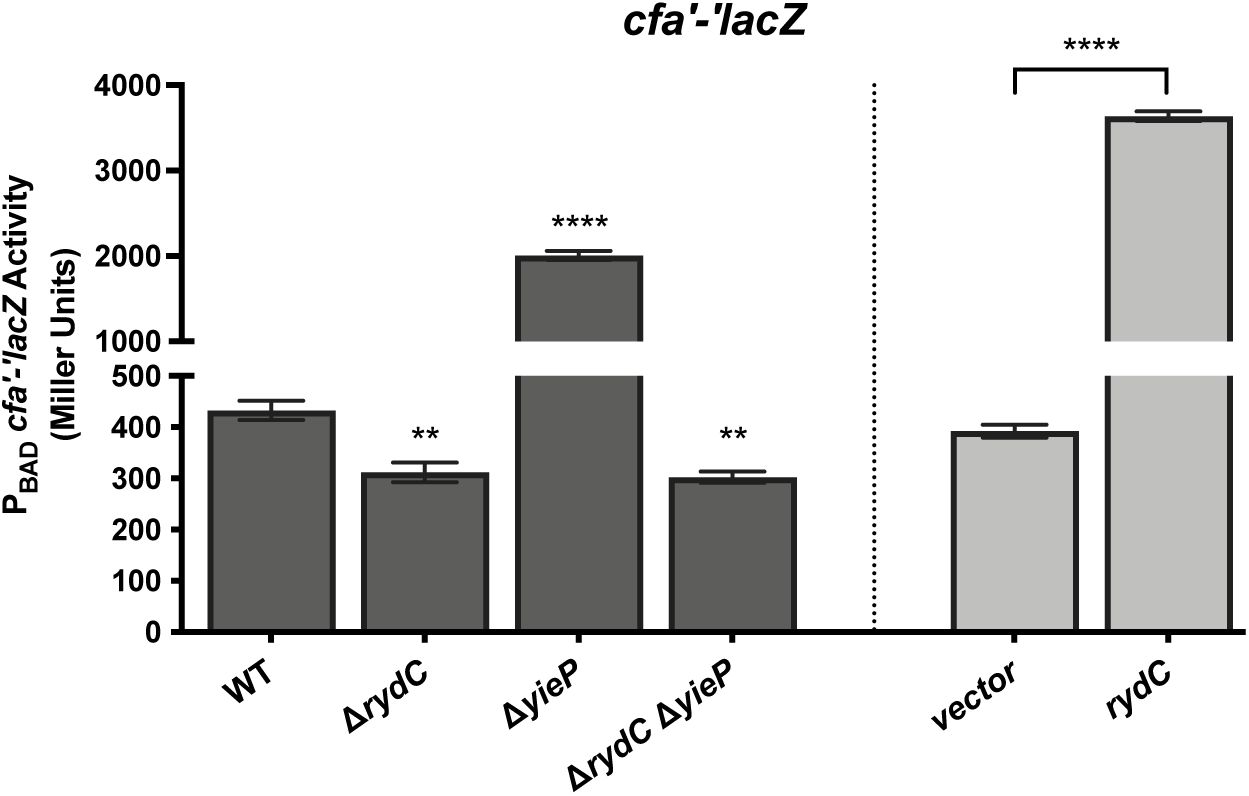
YieP alters *cfa* expression through RydC. Strains with a *cfa’-’lacZ* translational fusion in wild-type (WT), Δ*rydC*, Δ*yieP,* and Δ*rydC* Δ*yieP* mutant backgrounds (dark gray bars) were grown in TB medium with 0.002% L-arabinose to early exponential phase then samples were harvested and assayed for β-galactosidase activity. Error bars are standard deviations of the results of three independent experiments. Statistical significance was determined using a one-way ANOVA followed by Tukey’s post hoc test. Only significant differences between activity of WT compared to mutants is shown, (*, P < 0.05; **, P < 0.01; ***, P < 0.0001; ****, P < 0.0001). Separately, strains with a *cfa’-’lacZ* translational fusion carrying a vector control or P*_lac_*-*rydC* plasmid (light gray bars) were grown in TB medium with 0.002% L-arabinose to early exponential phase then *rydC* was induced with 0.1 mM IPTG. Samples were harvested 60 minutes later and assayed for β-galactosidase activity. The error bars represent standard deviations of three independent experiments. Expression of *rydC* significantly increases *cfa’-’lacZ* translational fusion activity compared to empty vector expression (two-tailed unpaired t-test, ****, P < 0.0001).

### Regulation of *yieP*

We next wanted to characterize the regulation of *yieP*. To study *yieP* transcription further, we first had to identify the transcription start site of *yieP*, as it has not been reported. We used template-switch-based 5’ Rapid Amplification of cDNA Ends (RACE) and determined the transcription start site is 27-nt upstream of the translational start site (Fig. 7A). Transcription factors commonly autoregulate their own transcription. To determine if YieP regulates its own expression, we constructed a *yieP’*-*lacZ* transcriptional fusion in the *yieP* locus and monitored activity in strains with a vector control or *yieP* expression plasmid (Fig. 7B). Expression of *yieP* led to a 2.4-fold increase in fusion activity, indicating that YieP can activate its own expression.

**Figure 7.**
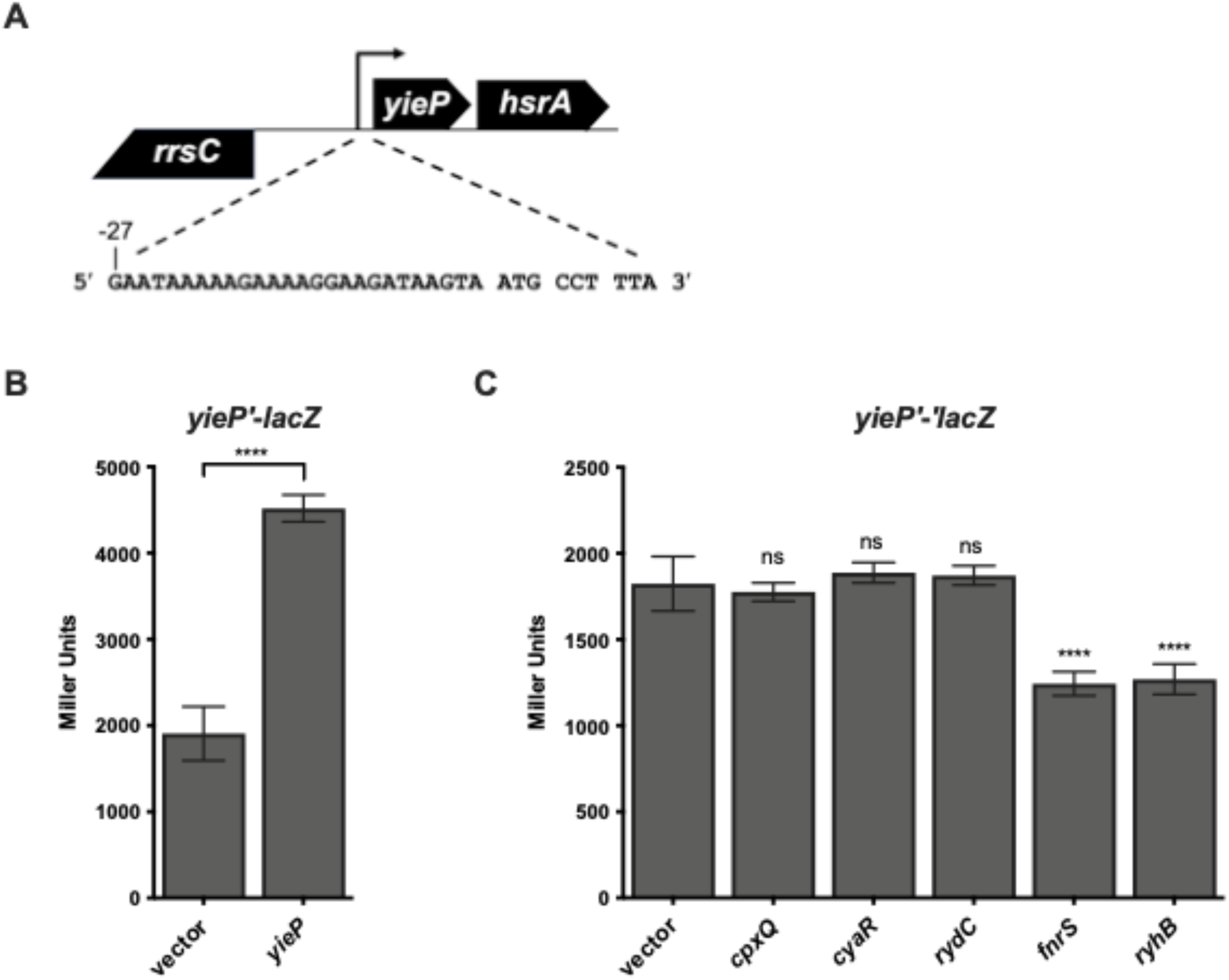
Regulation of *yieP* expression. (A) Template-switching 5’ RACE was used to determine the transcriptional start site of *yieP*. The sequence of the 27-nt 5’-UTR of *yieP* and the first three codons is shown below the gene diagram. (B) YieP autoinduces its own transcription. β-Galactosidase activity was measured in cells containing a *yieP*′-*lacZ* transcriptional fusion and a vector control or P_lac_-*yieP* plasmid. Strains harboring the *yieP’-lacZ* fusion with empty vector or *yieP*-expressing plasmid were grown overnight in TB containing ampicillin and IPTG. Cells were then subcultured the next day 1:100 into fresh TB with ampicillin and IPTG. Strains were grown at 37 °C with shaking for 3 hours and then β-galactosidase assays were performed. Expression of *yieP* significantly increases *yieP’-lacZ* fusion activity compared to empty vector expression (two-tailed unpaired t-test, ****, P < 0.0001). (C) Regulation of *yieP* translation by sRNAs. Strains with a *yieP’-’lacZ* translational fusion (controlled by the P_BAD_ promoter) and carrying a vector control or the indicated sRNA expression plasmids were induced with 0.002% L-arabinose and grown to early exponential phase. The sRNAs were induced by addition 0.1 mM IPTG and cells were grown for 1 hour, then samples were harvested and assayed for β-galactosidase activity. Error bars are standard deviations of the results of three independent experiments. Statistical significance was determined using a one-way ANOVA followed by Tukey’s post hoc test. Only significant differences between activity of the fusion with empty vector compared to when each sRNA expression is shown (ns, P not significant; *, P < 0.05; **, P < 0.01; ***, P < 0.0001; ****, P < 0.0001).

A recent RIL-seq (RNA interaction by sequencing and ligation) study uncovered potential *yieP* mRNA interactions with multiple sRNAs in *E. coli* [28]. To test whether *yieP* is also regulated post-transcriptionally by sRNAs, we made a *yieP’-’lacZ* translational fusion that included all putative sRNA interaction sites located in both the 5’ UTR and coding region (Fig. 7C). Ectopic production of candidate *yieP-*regulating sRNAs CpxQ and CyaR did not alter *yieP’-’lacZ* fusion activity while production of the sRNAs FnrS and RyhB led to a modest 1.5-fold reduction in activity (Fig. 7C). RydC was not predicted to interact with *yieP* mRNA using RIL-seq but we tested for RydC regulation of *yieP* because in some cases sRNAs and their transcription factors form regulatory feedback loops. However, expression of RydC did not alter *yieP* translation (Fig. 7C).

We chose to further characterize FnrS repression of *yieP* translation because we predicted that as a GntR type regulator, YieP might regulate genes involved in central metabolism and FnrS downregulates the levels of many mRNAs encoding enzymes involved in central metabolism [29]. Genetic analyses were conducted to confirm direct sRNA-mRNA base-pairing interactions. We first introduced a point mutation in FnrS (U61A for *fnrS-M*, Fig. 8A) that disrupted base pairing with *yieP* mRNA. The mutant sRNA, FnrS-M, could no longer repress *yieP’-’lacZ* (Fig. 8B). A compensatory mutation (A12U) in the *yieP* reporter fusion (*yieP’-’lacZ-*M) restored base-pairing interactions with FnrS-M (Fig. 8A). While wild-type FnrS could not repress the mutant *yieP’-’lacZ-*M fusion, FnrS-M repressed the *yieP’-’lacZ-*M fusion, with fold-repression restored to levels similar to the wild-type pair (Fig. 8C). These data are consistent with the model that FnrS base pairs with the translation initiation region of *yieP* mRNA, resulting in translational repression.

**Figure 8.**
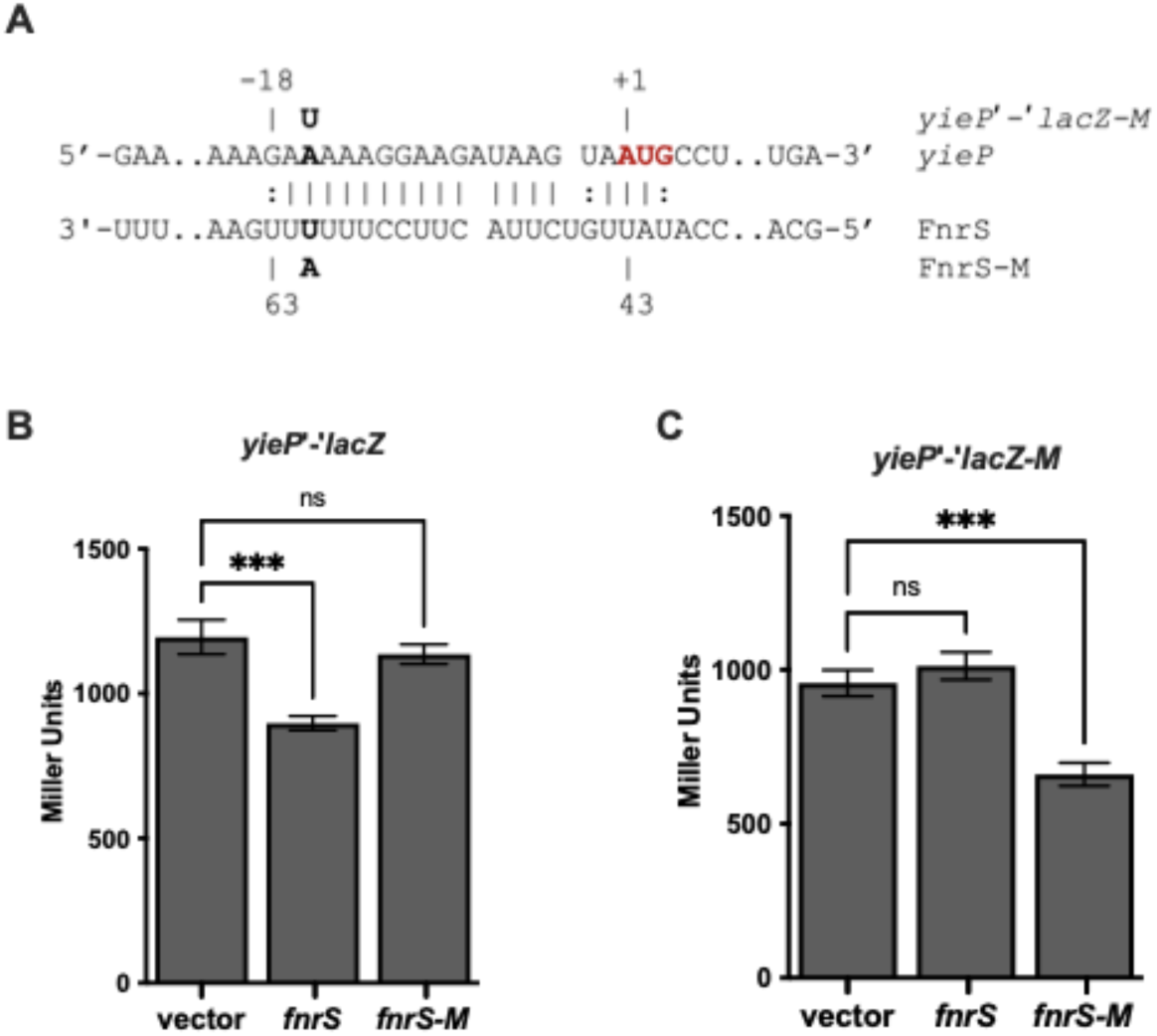
FnrS base pairs at the ribosome binding site of *yieP* mRNA to repress *yieP* translation. (A) Predicted base pairing between *yieP* mRNA and FnrS. The bolded nucleotides in *yieP* and FnrS were mutated to test the base-pairing prediction. The altered nucleotides in mutant constructs are shown above and below the base-pairing diagram. The start codon of *yieP* is indicated in red font. (B,C) Strains with wild-type *yieP’-’lacZ* or mutant *yieP’-’lacZ-M* fusions carrying vector control, FnrS, or mutant FnrS (FnrS-M) plasmids were grown with 0.002% L-arabinose to early exponential phase. The sRNAs were induced by addition 0.1 mM IPTG and cells were grown for 1 hour, then samples were harvested and assayed for β-galactosidase activity. The error bars represent standard deviations of three independent experiments. Statistical significance was determined using a one-way ANOVA followed by Tukey’s post hoc test. Only significant differences between activity of the fusion with empty vector compared to when each sRNA expression is shown (ns, P not significant; *, P < 0.05; **, P < 0.01; ***, P < 0.0001; ****, P < 0.0001).

### Regulation of RydC activity by a putative sponge sRNA

Most sRNAs have a transcriptional regulator like YieP that controls their production in response to certain growth or stress conditions. In addition, the activity of some sRNAs is controlled post-transcriptionally by so-called RNA sponges [30]. For example, SroC is an RNA produced by mRNA decay from the *gltIJKL* locus encoding an ABC transporter for amino acids. SroC acts as an RNA sponge for the sRNA GcvB and prevents GcvB from regulating its target, *gltIJKL* mRNA [30]. RIL-seq data identified RydC chimera with the sRNA OmrB, suggesting that RydC activity may be regulated by a sponge mechanism through pairing of RydC and OmrB [28]. OmrB production is controlled by the osmotic stress response regulator, OmpR [6]. To explore whether OmrB regulates RydC activity, we used the *cfa’-’lacZ* fusion as a reporter for RydC activity [18]. We first tested if ectopic production of OmrB altered *cfa’-’lacZ* activity in a WT background (Fig. 9A, WT). Expression of OmrB did not have an effect in this background, possibly because RydC is not produced at high levels under these conditions. Deletion of *yieP* results in increased RydC levels (Fig. 4C) which leads to increased activity of *cfa’-’lacZ* (Fig. 6 and Fig. 9A, Δ*yieP*). In the Δ*yieP* background, ectopic production of OmrB resulted in a 1.5-fold decrease in *cfa’-’lacZ* activity, indicating that OmrB may be able to inhibit RydC activity and regulation of *cfa* mRNA. To ensure that the OmrB-mediated decrease in *cfa* translation in the Δ*yieP* background was acting through RydC, OmrB was produced in a Δ*yieP* Δ*rydC cfa’-’lacZ* fusion strain (Fig. 9A). OmrB had no effect on the activity of the *cfa’-’lacZ* fusion in this background, supporting the model that OmrB acts indirectly on *cfa* by inhibiting RydC activity.

**Figure 9.**
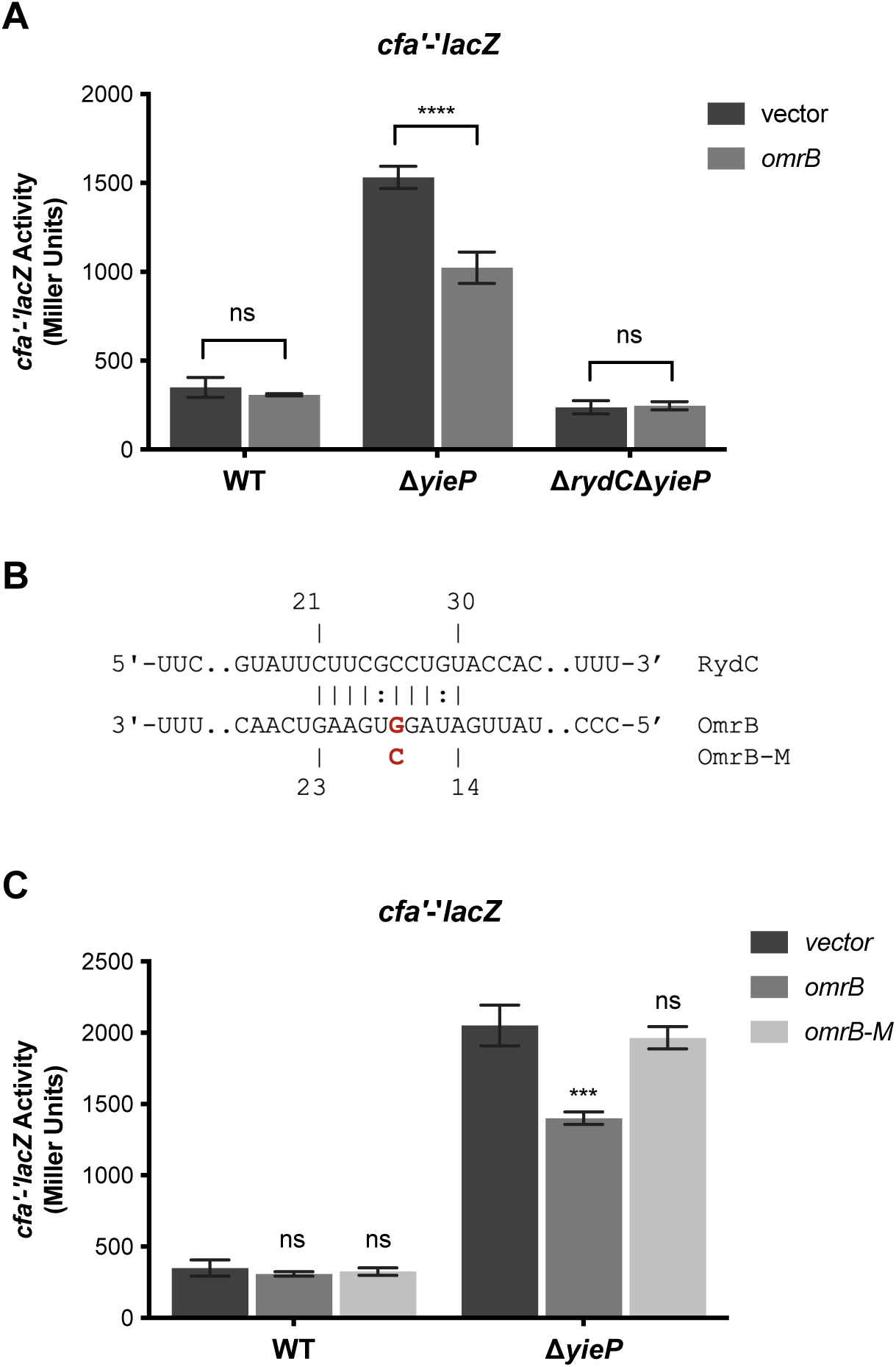
Regulation of RydC activity by OmrB sRNA. (A) Activity of the *cfa’-’lacZ* fusion in strains with the indicated genotypes carrying a vector control or *omrB* plasmid was measured. Strains were grown in TB medium with 0.002% L-arabinose and 0.1 mM IPTG to early exponential and samples were harvested and assayed for β-galactosidase activity. The error bars represent standard deviations of three independent experiments. Statistical significance of fusion activity in response to empty vector expression compared to *omrB* expression was determined using a two-tailed unpaired t-test, (ns, P not significant; *, P < 0.05; **, P < 0.01; ***, P < 0.0001; ****, P < 0.0001). (B) Predicted base pairing interaction between RydC and OmrB. One point mutation (G18C, OmrB-M) was made to disrupt base-pairing. (C) Activity of *cfa’-’lacZ* fusion in the indicated strains carrying a vector control, *omrB,* and *omrB-M* expression plasmids was measured as described in A. Statistical significance was determined using a one-way ANOVA followed by Tukey’s post hoc test. Only significant differences between activity of the fusion with empty vector compared to when each sRNA expression is shown, (ns, P not significant; *, P < 0.05; **, P < 0.01; ***, P < 0.0001; ****, P < 0.0001).

We propose that OmrB prevents RydC from regulating the *cfa’-’lacZ* translational fusion by base pairing with RydC and making it unavailable for interaction with *cfa* mRNA. Ectopic production of OmrB did not alter activity of *rydC*′-*lacZ* compared to vector control, indicating that OmrB does not indirectly increase *rydC* transcription (Fig. S3). A predicted OmrB-RydC base-pairing interaction is shown in Fig. 9B. We made a point mutation in OmrB, yielding OmrB-M, that should weaken the base-pairing interaction. OmrB-M had no impact on *cfa’-’lacZ* activity in a wild-type background and in the Δ*yieP* mutant background, OmrB-M lost the repressive effect demonstrated by OmrB (Fig. 9C). These data are consistent with the idea that OmrB serves as a sponge for RydC to antagonize RydC activity under osmotic stress conditions, when OmrB is produced at high levels.

## Discussion

Regulation of membrane lipid composition is an important adaptation to stress [1, 31, 32] and control of CFA levels through transcriptional [33, 34] and post-transcriptional regulators responding to different stresses suggests that incorporation of CFAs into membrane lipids is important for adaptation to a variety of stressful environments. In this study, we tried to understand more about the physiological responses that modulate *cfa* mRNA levels through an important activating sRNA, RydC. We found that both RydC and *cfa* promote resistance to membrane-disrupting stresses including exposure to butanol and SDS/EDTA (Fig. 2). Despite clear phenotypes, we found no evidence that *rydC* transcription is regulated by these same membrane-disrupting conditions or through known membrane stress regulators (Figs. 3 and S2). Instead, we find that *rydC* transcription is repressed by a GntR-family transcription factor, YieP (Figs. 4,10). We also discovered additional sRNA regulators of this pathway that contribute indirectly to *cfa* control. FnrS represses *yieP* translation (Fig. 7), and thus likely indirectly activates *cfa* via derepression of RydC (Fig. 10). OmrB antagonizes RydC-mediated activation of *cfa* (Fig. 9), likely through a sponge-like mechanism (Fig. 10). Collectively, our results are consistent with the model shown in Figure 10, where multiple stress response pathways responding to different signals converge to regulate *cfa* directly and indirectly through transcriptional and post-transcriptional mechanisms.

**Figure 10.**
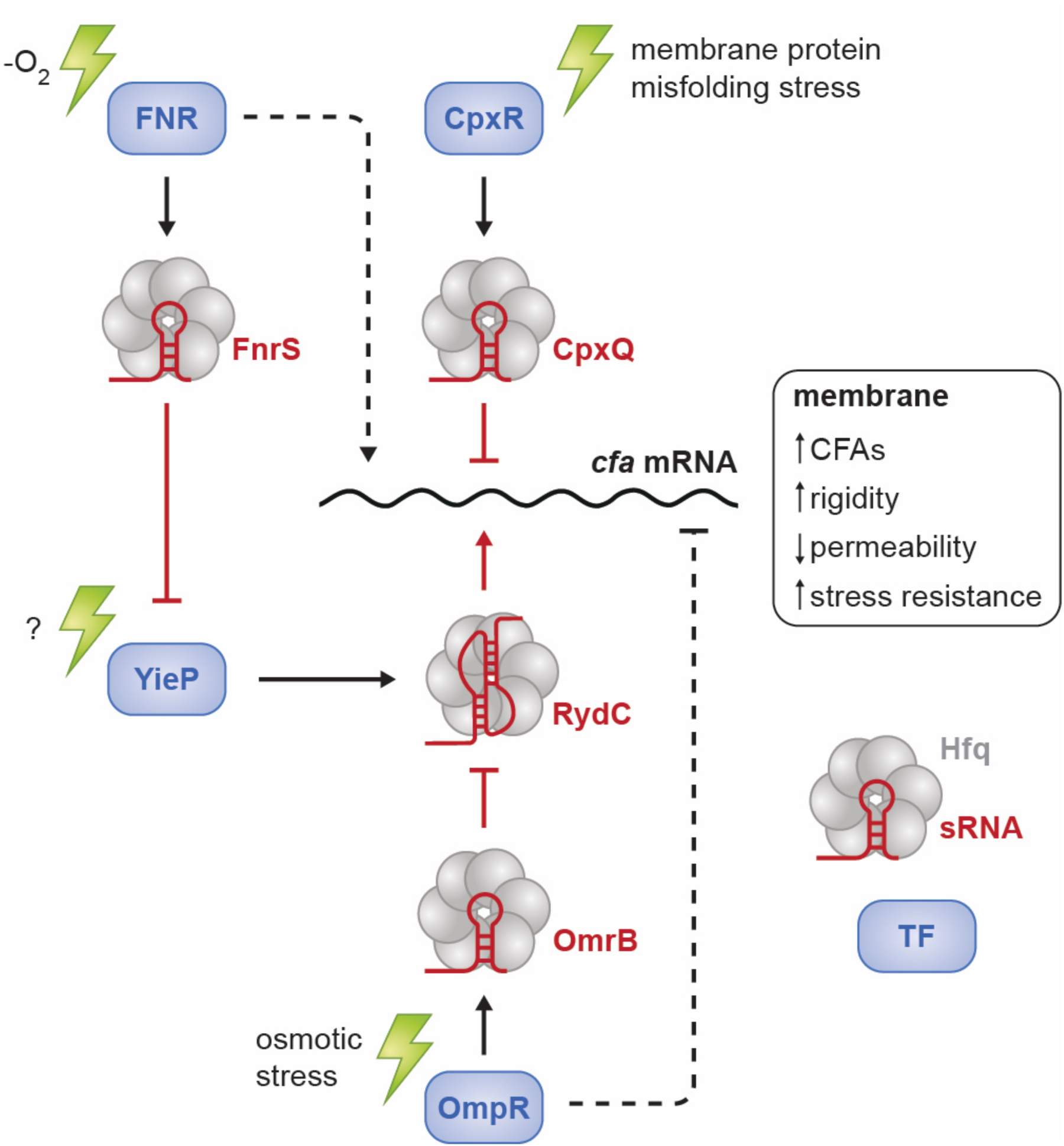
Transcriptional and post-transcriptional regulatory network controlling production of cyclopropane fatty acids. Transcription factors and sRNAs that are directly or indirectly involved in regulation of *cfa* mRNA translation or stability are illustrated. Transcription factors are indicated in blue. sRNAs are indicated in red. Stress signals promoting the indicated response are illustrated by green symbols. Arrows indicate activation, blunt lines indicate repression. Solid lines indicate direct regulation, dotted lines indicate indirect regulation.

Transcription of *cfa* itself is controlled by two different promoters – a σ^70^ promoter that produces the *cfa* long isoform that is subject to regulation by sRNAs [11, 12], and a σ^S^ promoter that produces a short sRNA-insensitive isoform during stationary phase [33]. Beyond this, at minimum an additional four transcription factors, FNR, YieP, OmpR, and CpxR regulate *cfa* indirectly through their control of sRNAs (Fig. 10). Direct regulation of *cfa* by sRNAs has been demonstrated previously [11, 12, 17]. RydC activates *cfa* by antagonizing RNase E and Rho activity within the *cfa* 5’ UTR while CpxQ represses *cfa* by a mechanism that also involves RNase E and Rho [11, 12, 17]. We now know that RydC levels are controlled by the repressor YieP, but the stress signal sensed by YieP remains a mystery. Interestingly, while RydC promotes resistance to membrane disrupting conditions through its regulation of *cfa* (Fig. 2), these and other membrane stress conditions and regulators do not control RydC transcription (Figs. 3, S2). This implies that basal levels of RydC are sufficient to promote a physiologically important level of activation of CFA synthesis. CpxQ is controlled by CpxR, the response regulator component of the Cpx pathway that responds to a variety of signals that cause membrane protein misfolding [2, 10]. In this study, we find evidence for additional indirect sRNA regulatory inputs in control of *cfa* expression. FnrS, an sRNA that participates in regulation of multiple mRNAs under anaerobic growth conditions, negatively regulates translation of *yieP*, encoding the repressor of *rydC* transcription. We expect that by repressing a repressor, FnrS promotes RydC production and *cfa* activation. OmrB is another indirect sRNA regulator of *cfa* that we identified in this study. Production of this sRNA is activated by OmpR under osmotic stress conditions [6] and we find evidence for OmrB antagonism of RydC activity. By repressing an activator, we expect that OmpR/OmrB indirectly inhibit *cfa* expression during the osmotic stress response.

While we do not fully understand the physiological significance of *cfa* regulation by these sRNAs or how these responses are adaptive during stress, it is noteworthy that all of the relevant signals and regulatory pathways are linked in some way to membrane stress. The Cpx two-component system senses conditions that induce membrane protein misfolding [10, 35] which leads to CpxR-mediated activation of genes encoding proteases and protein folding chaperones that help to restore homeostasis [22, 36–40]. CpxR also represses transcription of genes encoding high molecular weight protein complexes including those involved in the aerobic respiratory electron transport chain, presumably to avoid overaccumulation of large unfolded or mis-folded protein complexes that would disrupt membrane structure and function [23, 41]. In this study, we found that FNR, a global regulator of anaerobic metabolism [29] potentially indirectly activates *cfa* through the sRNA FnrS, which directly regulates *yieP* translation (Figs. 8, 10). FNR and FnrS repress genes involved in aerobic metabolism and activate genes involved in anaerobic metabolism. Because the shift from aerobic to anaerobic metabolism involves substantial changes in the need for specific membrane respiratory complexes, we hypothesize that altering CFA levels in membrane lipids may play a role in allowing cells to maintain membrane protein structure and function during this shift.

This study also revealed a novel regulatory link between the osmotic stress response and modulation of CFAs in *E. coli*. High osmotic stress increases CFAs in lactic acid bacteria [42] and halotolerant bacteria naturally have high levels of CFAs in their membranes [43]. Heterologous overproduction of a CFA synthase from a halophilic organism increased *E. coli* tolerance to various stress conditions, including high salt [44]. In *E. coli,* the response to osmotic stress is mediated by the OmpR-EnvZ two-component system [45, 46]. Osmotic stress also induces RpoS [47–49], which would increase *cfa* transcription from the proximal promoter [33]. The OmpR-activated sRNA OmrB post-transcriptionally downregulates *cfa* under conditions where RydC is produced at high levels. These opposing effects, and the ultimate outcome for CFA levels is likely highly dependent on the specific combination of environmental signals present.

A key remaining question is what signal triggers YieP derepression of *rydC,* leading to strong upregulation of CFA production. YieP belongs to the GntR family of transcription factors that was first described in 1991 and named after the gluconate operon repressor of *Bacillus subtilis* [50]. The GntR family is one of the largest families of bacterial transcription factors and is found across many bacterial species where it has been shown to regulate genes involved in carbohydrate transport and metabolism [51, 52]. GntR-family regulatory proteins consist of two domains, an N-terminal DNA-binding domain and C-terminal ligand-binding domain [53]. The DNA-binding domain is a helix-turn-helix (HTH) motif and is similar among all GntR-family transcription factors while the larger C-terminal ligand-binding domain is less conserved among GntR-family transcription factors [51, 53]. Most GntR family members act as homodimers, with oligomerization occurring at the C-terminus. GntR family members typically recognize palindromic sequences of DNA that are A/T rich [54]. As we observed for YieP regulation of *rydC,* most GntR-type transcription factors act as repressors of transcription [55]. Upon ligand binding, the GntR-type transcription factor is removed from the operator site and transcription of downstream genes occurs [55]. YieP acts as an activator of its own transcription, and presumably also activates transcription of the downstream efflux pump gene *hsrA*. Further work is needed to determine if YieP repression and activation activities both require binding to a small molecule.

A phylogenetic tree made from the full-length multiple alignment of the C-terminal domain of GntR members revealed four types or subfamilies [51]. Based on its C-terminal domain, YieP belongs to the FadR-type subfamily, which is named after the repressor (FadR) of fatty acid degradation in *E. coli* [56]. The FadR subfamily is the largest subfamily and contains 40% of GntR regulators [51]. Most of the FadR-family proteins are involved in the regulation of oxidized substrates related to amino acid metabolism or the intersection of various metabolic pathways [51]. GntR regulators in this subfamily bind ligands such as: pyruvate (PdhR, *E. coli*), glycolate (GlcC, *E. coli*), galactonate (DgoR, *E. coli*), lactate (LldR, *E. coli*), gluconate (GntR, *B. subtilis*), aspartate (AnsR, *Rhizobium etli*), and malonate (MatR, *Rhizobium leguminosarum*) [51]. A putative regulon of YieP has been determined by chromatin immunoprecipitation with sequencing combined with exonuclease treatment (ChIP-exo), which detects *in vivo* interactions between transcription factors and target genes at base-pair resolution [57]. YieP’s putative regulon includes the uncharacterized genes *ykgE* and *yohJ*, as well as *arcB* (anoxic redox control histidine kinase), *dauA* (succinate, fumarate, and aspartate transporter), and *glpD* (glycerol 3-phosphate dehydrogenase) [57]. YieP was found to commonly bind downstream of the promoter of these genes, suggesting YieP acts primarily as a repressor [57]. Although the regulon of YieP has been identified, ChIP-exo was only able to identify locations that YieP binds under the condition tested (exponential growth in glucose minimal medium) and did not validate or characterize the effects of this binding [57]. Moreover, this putative regulon did not include *rydC*, whose transcription we show is repressed by YieP.

Phenotypes associated with loss of RydC and *cfa,* along with the known functions of sRNAs and proteins involved in this regulatory network paint a picture of *cfa* as a hub for membrane stress signal integration. Further work is needed to better understand the specific contributions of protein and sRNA regulators to *cfa* control and the biophysical properties of CFA-containing membranes under a variety of conditions.

## Materials and Methods

### Strain and plasmid construction

Strains and plasmids used in this study are listed in Table S1. All strains used in this study are derivatives of *E. coli* K12 DJ624 (D. Jin, National Cancer Institute) or *Salmonella enterica* serovar Typhimurium ATCC 14028 (American Type Culture Collection). Oligonucleotide primers used in this study were synthesized by Integrated DNA Technologies and are listed in Table S2.

### *E. coli* gene deletions and plasmids

All gene deletions in *E. coli* were made via P1 *vir* transduction from the Keio collection [58] into a clean background (either DJ624 or DJ480). The kanamycin marker was removed using the FLP recombinase, encoded on the temperature-sensitive pCP20 plasmid [59]. Gene deletions were confirmed with PCR and Sanger sequencing.

Plasmids containing *yieP*, *rydC*, *cpxQ*, *cyaR*, *fnrS*, or *ryhB* under the control of the P_Llac_ promoter were constructed by PCR amplifying the gene from *E. coli* DJ624 chromosomal DNA using oligonucleotides containing HindIII and BamHI restriction sites (Table S2). PCR products and vector pBRCS12 were digested with HindIII and BamHI (New England Biolabs) restriction endonucleases. Digestion products were ligated using DNA Ligase (New England Biolabs) and the plasmids were transformed into XL10 competent cells and the plasmid was purified. All plasmids were confirmed by Sanger sequencing. Plasmids harboring mutated sRNA alleles under the control of the P_Llac_ promoter were constructed using the QuikChange II XL site-directed mutagenesis kit (Agilent Technologies, Santa Clara, CA) with oligonucleotides that contain mismatched bases at the desired locations (Table S2) and transformed into XL10-Gold Ultracompetent cells.

### E. coli lacZ fusions

Two *rydC* transcriptional fusions were made: an in-locus *lacZ* fusion and an out-of-locus *lacZ* fusion. The in-locus *lacZ* fusion to *rydC* (CB227) was constructed as described previously [60]. Briefly, the primers CB4F and CB4R, which contain 40 nt of sequence homologous upstream and downstream of *rydC* and 20 nt of sequence homologous to the FLP recombination target (FRT)-chloramphenicol cassette, were amplified using pKD3 as the template. This DNA fragment was recombined into the bacterial chromosome at the *rydC* locus using λ red recombination. Using PCP20, the chloramphenicol cassette was flipped out and *lacZ* was inserted using pKG136 plasmid (derivative of pCE36). Using P1 *vir*, the in-locus *rydC’*-*lacZ* fusion was moved into *E. coli* WT and deletion backgrounds.

The out-of-locus *rydC* transcriptional fusion (CB121) was constructed using λ Red homologous recombination into strain PM1205 and counter-selection against *sacB*, as previously described [61]. Primers CB7 and CB8 were designed with homology to the region upstream of P_BAD_ and *lacZ*, to ensure P_BAD_ was replaced upon recombination. The fusion contains the 171 nucleotides upstream of *rydC* and the first 5 nucleotides of RydC.

The *cfa’-’lacZ* translational reporter fusion [18] was constructed using λ Red homologous recombination into strain PM1205 and counter-selection against *sacB,* as previously described [61]. Transcription of the fusion is under control of the P_BAD_ promoter and the fusion contains the full 212-nt *cfa* 5′ UTR and the first 36 codons of the *cfa* open reading frame.

Both *yieP’*-*’lacZ* translational reporter fusions (*yieP’-’lacZ* and *yieP’-’lacZ-*M) were constructed using λ Red homologous recombination into strain PM1205 and counter-selection against *sacB* [61]. Transcription of the fusion is under control of the P_BAD_ promoter and the fusion contains the entire 27-nt of the *yieP* 5’ UTR fragment of interest and the entire open reading frame except the stop codon. Primers CBY1 and CBY2 were used to generate dsDNA substrate for *yieP’-’lacZ*. CBY3 and CBY2 were used to generate dsDNA substrate for *yieP’-’lacZ-*M, with CBY3 containing the point mutation of interest.

Two *yieP* transcriptional fusions were made: an in-locus *lacZ* fusion and an out-of-locus *lacZ* fusion. The in-locus lacZ fusion to the *yieP* promoter (CB308) was constructed as described above for the in-locus *lacZ* fusion to *rydC* and previously [60]. However, the primers CB47 and CB48 were used to amplify pKD3. P1 vir, was used to move the in-locus *yieP’*-*lacZ* fusion into *E. coli* DJ624.

The out-of-locus *yieP* transcriptional fusion (CB331) was constructed using λ Red homologous recombination into strain PM1205 and counter-selection against *sacB*, as previously described [61]. Primers CB66 and CB67 were designed with homology to the region upstream of P_BAD_ and *lacZ*, to ensure P_BAD_ was replaced upon recombination. The fusion contains the 481 nucleotides upstream of *yieP* and ends right before the start codon.

### Salmonella constructs

The *rydC* transcriptional fusion was constructed as described previously [60]. Briefly, the primers CB78 and CB86, which contain 40 nt of sequence homologous upstream and downstream of *rydC* and 20 nt of sequence homologous to the FLP recombination target (FRT)-chloramphenicol cassette, were amplified using pKD3 as the template. This DNA fragment was recombined into the bacterial chromosome at the *rydC* site using λ red recombination. Using pCP20, the chloramphenicol cassette was flipped out and *lacZ* was inserted using pKG136 plasmid (derivative of pCE36). The fusion was verified via PCR [60] and then transduced into clean WT and Δ*yieP* backgrounds using phage P22 HT105/1 int-201 [62].

The Δ*yieP* mutation in *Salmonella* was constructed using λ Red-mediated recombination as previously described [59, 60] resulting in insertion of a kanamycin resistance cassette. Briefly, primers CB72 and CB73 were used to amplify the kanamycin cassette in pKD13. The PCR product was transformed into *S. enterica* carrying pKD46, the plasmid for RED-recombineering. The deletion was verified via PCR and then transduced into a clean background using phage P22 HT105/1 int-201 [62]. The kanamycin resistance cassette was removed using the pCP20 plasmid [59].

Plasmids containing *yieP* under the control of the P_Llac_ promoter were constructed by PCR amplifying the gene from *Salmonella* chromosomal DNA using the above protocol for *E. coli* and were constructed in *E. coli* Top10. Plasmids containing P_lac_-*yieP* and empty vector control were passaged through a restriction-minus modification-plus Pi1 *Salmonella* strain JS198 [60] prior to transformation into *Salmonella enterica* serovar Typhimurium ATCC 14028 backgrounds. Plasmids were verified using Sanger sequencing,

### Media and growth conditions

Bacteria were cultured in LB broth medium or on LB agar plates at 37°C, unless otherwise stated. When necessary, media were supplemented with antibiotics at the following concentrations: 100 μg ml^−1^ ampicillin (Amp), 25 μg ml^−1^ chloramphenicol (Cm), or 25 μg ml^−1^ kanamycin (Kan). Isopropyl β-D-1-thiogalactopyranoside (IPTG) was used at 0.1 mM (final concentration) for induction of expression from the P_lac_ promoter. For plates, LB agar was autoclaved and cooled to 56°C. Various stressors were added to the following concentrations: 0.5% SDS and 0.5 mM EDTA or 1% butanol. All plates contained ampicillin and IPTG. Strains were streaked the same day plates were made and incubated overnight at 37°C. **Transposon screen for regulators of *rydC* transcription**

Strains carrying a *rydC’-lacZ* transcriptional fusion were electroporated with the Tn5-mini transposon pRL27 [63] and screened on MacConkey agar with lactose and kanamycin. Red colonies were picked and purified twice to confirm phenotype. Approximately 6000 colonies were analyzed, with only two red colonies found, both with Tn insertions in *yieP*. Locations of transposon insertions were determined using plasmid recovery, as described previously [64].

### Template switching 5′ RACE

RNA was extracted from DJ624 using the hot phenol method [65]. 5′ RACE was performed using the “two-step” method of Pinto and Lindblad [66]. Briefly, RNA denaturation took place at 65 °C, for 5 min, in the presence of dNTPs. The temperature was then lowered to 50 °C and the cDNA elongation mastermix was added to the denatured RNA. Synthesis of cDNA was primed by a gene-specific primer located in the coding region of *yieP* (GSP) using SuperScript III. SuperScript III adds extra cytosines to the 5′ end of the mRNA, resulting in cDNA with a poly-C. After incubation at 50 °C for 60 min, the template-switching mastermix was added, the temperature was lowered to 42 °C, and the reactions were incubated for an additional 90 min. The template-switching mastermix contains a template switching oligo (TSO) that contains a 3′ poly-G tail which pairs with the cDNA poly-C and is used as template for reverse transcriptase. To inactivate reverse transcriptase, the reaction was put at 85 °C for 5 min then cooled. RNase H was added and the reaction incubated a 37 °C for 20 min to degrade leftover RNA. Then, two rounds of PCR were completed. The first-round PCR was carried out using a universal outer primer (OUTER) and the gene specific primer used in the initial reverse transcription reaction (GSP) with the cDNA as template. The OUTER primer contains a part of the TSO. The resulting product was then diluted 100-fold before being used as the template for the second-round PCR, which was performed using a universal inner primer (INNER) and a gene specific primer (GSP2). INNER is identical to the 3′ end of the OUTER primer. All PCR reactions were performed using Q5 High-Fidelity 2X Master Mix (NEB, Frankfurt, Germany). The PCR product was purified and sequencing was done using a gene specific primer (GSP3). The final location of the transcription start site was determined based on three independent 5′ RACE reactions.

### β-Galactosidase assays

Bacterial strains harboring *rydC’-lacZ* reporter fusions in mutant backgrounds (Fig. 3) were cultured overnight in tryptone broth (TB) and then subcultured the next day 1:100 into fresh TB. Strains were grown at 37 °C with shaking for 3 hours and then β-galactosidase assays were performed as previously described [67].

*E. coli* and *Salmonella* strains harboring the *rydC’-lacZ* and carrying either vector or P_lac_-*yieP* plasmids (Fig. 4) were cultured overnight in TB with Amp and 0.1 mM IPTG and then subcultured 1:100 to fresh TB medium containing Amp and 0.1 mM IPTG. Strains were grown at 37 °C with shaking for 3 hours and then β-galactosidase assays were performed as previously described [67].

Strains harboring the *cfa’-’lacZ* translational reporter fusion with or without vector control and sRNA-expressing plasmids (Figs. 6,9) were cultured overnight in TB with Amp and 0.002% L-arabinose then subcultured 1:100 to fresh TB medium containing Amp and 0.002% L-arabinose. Strains were grown at 37°C with shaking to early exponential phase then 0.1mM IPTG was added to induce sRNA expression. Samples were harvested 60 minutes later and β-galactosidase assays were then performed as previously described [67].

Strains harboring the out of locus *yieP’-lacZ* transcriptional fusion with empty vector or *yieP*-expressing plasmid (Fig. 7B) were grown overnight in TB containing amp and IPTG. Cells were then subcultured the next day 1:100 into fresh TB with amp and IPTG. Strains were grown at 37 °C with shaking for 3 hours and then β-galactosidase assays were performed as previously described [67].

Strains harboring *yieP’-’lacZ* translational fusion with vector or sRNA-expressing plasmids (Figs. 7C, 8) were grown as described above for *cfa’-’lacZ* translational reporter fusion.

For all β-galactosidase assays, Miller units are defined as (micromoles of *o*-nitrophenol [ONP] formed per minute) × 10^6^/(optical density at 600 nm [OD_600_] × milliliters of cell suspension).

### Northern blot

To measure RydC levels by Northern blot (Fig. 4C), bacterial strains were grown overnight then subcultured 1:100 to fresh LB medium (four biological replicates). Subcultures were grown for 2 hours and then RNA was extracted using the hot phenol method as described previously [65]. For the positive control strain producing RydC from the inducible P_lac_-*rydC* plasmid, cells were grown overnight in LB Amp, then subcultured 1:100 to fresh LB Amp medium. Subcultured cells were grown for 2 hours, 0.1 mM IPTG (final concentration) was added to induce *rydC,* and RNA was extracted 20 minutes later. RNA concentrations were measured spectrophotometrically.15 μg of RNA were denatured for 5 min at 95°C in loading buffer (containing 95% formamide), separated on 8% polyacrylamide urea gel at 100 V for 1 h using 1× Tris-acetate-EDTA (TAE), then transferred to BrightStar™-Plus Positively Charged Nylon Membrane in 0.5× TAE buffer by electroblotting at 50 V for 1 hour at 4°C. RNA was crosslinked to the membrane, then the membrane was prehybridized for 45 min in ULTRAhyb (Ambion) solution at 42 °C. Blots were hybridized overnight with ^32^P-end-labeled primer “RydC NB,” then membranes were washed at 42 °C twice with 2X SSC/0.1% SDS for 8 minutes then twice with 0.1X SSC/0.1% SDS for 15 minutes. The signal was detected using film. Membranes were stripped in boiling 0.1% SDS for 10 minutes then re-probed overnight ^32^P-end-labeled primer “5S.” The ladder was prepared using the Decade Markers System from ThermoFisher, according to manufacturer’s instructions.

## Acknowledgements

We would like to thank John Cronan, James Imlay, James Slauch, and members of the Slauch lab for thoughtful discussions and advice. We also thank members of the Vanderpool lab for strains, plasmids, advice, and discussions. We are grateful to Dr. Sandy Westermann for graphic design. This work was supported by the National Institutes of Health R35 grant (R35 GM139557) awarded to C.K.V.

